# A *trans*-piRNA network and transcriptional antagonism shape piRNA cluster function

**DOI:** 10.64898/2026.04.08.717276

**Authors:** Yicheng Luo, Daniel Siqueira de Oliveira, Peng He, Alexei A. Aravin

## Abstract

PIWI-interacting (pi)RNAs protect animal germlines from transposable elements (TEs) by guiding their sequence-specific repression. In *Drosophila* germline, piRNAs are encoded in distinct genomic regions, piRNA clusters (piCs), that are transcribed by a non-canonical machinery that is anchored on chromatin by the HP1 paralogue Rhino. Studies of transgenic piCs revealed that piRNA biogenesis depends on cytoplasmic inheritance of piRNAs, however, whether native piCs require trans-generational piRNA transmission remained unknown. Here, we used two approaches to show that cytoplasmic inheritance of cognate piRNAs is critical for piRNA biogenesis. Our analyses reveal that individual piCs form a tightly interconnected network linked by *trans*-acting piRNAs that reinforce biogenesis. According to the transposon trap model, the content of piCs is updated by integration of novel TEs leading to production of piRNA guides against integrated transposons. However, we found that transcription driven by promoters integrated into piCs disrupt local piRNA biogenesis by removing Rhino and antagonizing non-canonical transcription of piRNA precursors. Thus, newly inserted transposons might suppress piRNA production before they become domesticated by the piRNA pathway calling for a revision of the trap model. Together, our results reveal that piC activity is shaped by transcriptional competition and a dynamic interplay between individual piCs connected into a common network.

**Highlights:**

- Cytoplasmic inheritance of maternal piRNAs is required for piRNA biogenesis in the next generation
- *Trans*-acting piRNA ensure robustness of piRNA biogenesis and repression by connecting individual clusters into a functional network
- Ping-pong amplification provides non-uniform processing of different regions inside piCs
- Genes can remain active inside piRNA clusters antagonizing Rhino-dependent non-canonical transcription and piRNA biogenesis, challenging the transposon trap model

## Introduction

Transposable elements (TEs) are pervasive genomic parasites that threaten genome integrity in animal germlines. To counteract TE activity, metazoans evolved the PIWI-interacting RNA (piRNA) pathway, a small RNA–guided defense mechanism that detects and silences transposon transcripts^1^. piRNAs associate with PIWI-clade Argonaute proteins and direct repression of complementary RNA targets through both post-transcriptional and transcriptional mechanisms. In *Drosophila*, two Piwi-clade proteins, AUB and Ago3, cleave TE transcripts in the cytoplasm, generating secondary piRNAs through the ping-pong amplification cycle^2^. Nuclear PIWI protein recognizes nascent transcripts and induces transcriptional silencing associated with deposition of the H3K9me3 chromatin mark^3–6^.

In *Drosophila melanogaster*, germline piRNAs originate from genomic loci known as piRNA clusters (piCs)^2^ which harbor fragmented remnants of TEs and function as genomic archives of past TE invasions. piCs generate long precursor transcripts that are processed into mature piRNAs, thereby producing sequence information that guides recognition of active TEs elsewhere in the genome. Transcription of dual-strand clusters is driven by a specialized non-canonical transcription (NCT) machinery centered on the HP1 homolog Rhino (Rhi), which binds H3K9me3-enriched chromatin and recruits additional cofactors^7–12^. Rhi and its partners promote promoter-independent transcription of piRNA precursors while suppressing canonical mRNA processing events and were also proposed to direct transcripts to the cytoplasmic processing machinery^7,13–15^.

Previous studies suggested that maternally inherited piRNAs provide a trigger for piC activation in the next generation. During oogenesis, piRNAs produced in the maternal germline are deposited into the oocyte and preserved in the early embryo, where they initiate piRNA production. Experiments using artificial transgenic piCs demonstrated that maternally inherited piRNAs convert cognate sequences into active piRNA-producing loci accompanied by deposition of the H3K9me3 mark and recruitment of Rhi^10,16–18^. However, whether maternally inherited piRNAs are required to maintain piRNA biogenesis at endogenous clusters remains unclear.

Adaptation of the piRNA pathway to the newly invading TEs presents another twist to the complex relationship between piCs and TEs. Unlike prokaryotic CRISPR defense systems, the piRNA pathway lacks dedicated machinery that would insert fragments of TE sequences into piCs to produce novel guides for repression. Instead, piCs seem to rely on the intrinsic activity of TEs themselves. According to the ‘trap model’, active TEs occasionally integrate into piCs, where they become sources of antisense piRNAs that silence homologous elements elsewhere in the genome. Insertions of TE fragments accumulate within clusters, forming repositories of sequences that provide immunity against TE activity^19–23^. The trap model is supported by observations that heterologous sequences inserted into piCs produce piRNA^17,19,24^. The trap model suggests that TE inserted into piRNA clusters are rapidly silenced and converted into sources of piRNAs. However, whether newly inserted TEs retain transcriptional activity and how such activity interacts with the specialized transcriptional environment of piCs remain open questions.

Here, we explore the role of cytoplasmic inheritance of piRNAs and investigate how transcriptional dynamics influence the trapping of active genes in piCs. Using deletions as well as engineered insertions into native piRNA clusters, we demonstrate that maternal piRNAs are essential for jump-starting piRNA biogenesis in the next generation. Our results reveal that piCs operate within an extensive network interconnected by *trans*-acting piRNAs. We show that canonical transcription (CT) of a gene inserted into a piC interferes with non-canonical transcription of precursors and with piRNA production by local displacement of Rhi from chromatin, suggesting that the domestication of TEs thorugh their integration into piCs involves a competition between transcriptional programs. Together, our work provides a revised framework for understanding functions of piRNAs in maintaining genome defense and adaptation to invading genetic elements.

## Results

### Impact of maternal piRNAs on piRNA biogenesis

To determine if cytoplasmic trans-generational piRNA inheritance is required for the function of endogenous piCs we analyzed piRNA biogenesis in progenies of two reciprocal crosses between wild-type flies and flies with deletion of individual piCs (Fig. 1A, left panel). Such progenies are genetically identical, but different in respect to cytoplasmic inheritance of cluster piRNA: the progeny from cross of wild type mothers and fathers with cluster deletion (termed maternal cross) but not the reciprocal (paternal) cross receives cluster piRNAs from their mothers. Surprisingly, cytoplasmic inheritance of maternal *42AB*-derived piRNAs had little impact on piRNA biogenesis in the progeny: progeny that lacked maternal *42AB* piRNAs had only 14.5% reduction in the level of *42AB* piRNA (n=2, p=0.018, two-tailed Student’s test) (Fig. 1B). For *38C* cluster deletion, reduction of piRNA biogenesis upon absence of maternal *38C* piRNAs was even less pronounced at 5.9% (n=2, p=0.119) (Fig. 1B). Thus, maternal piRNAs generated by *42AB* and *38C* clusters are largely dispensable for the activity of these clusters in the progeny.

**Figure 1.**
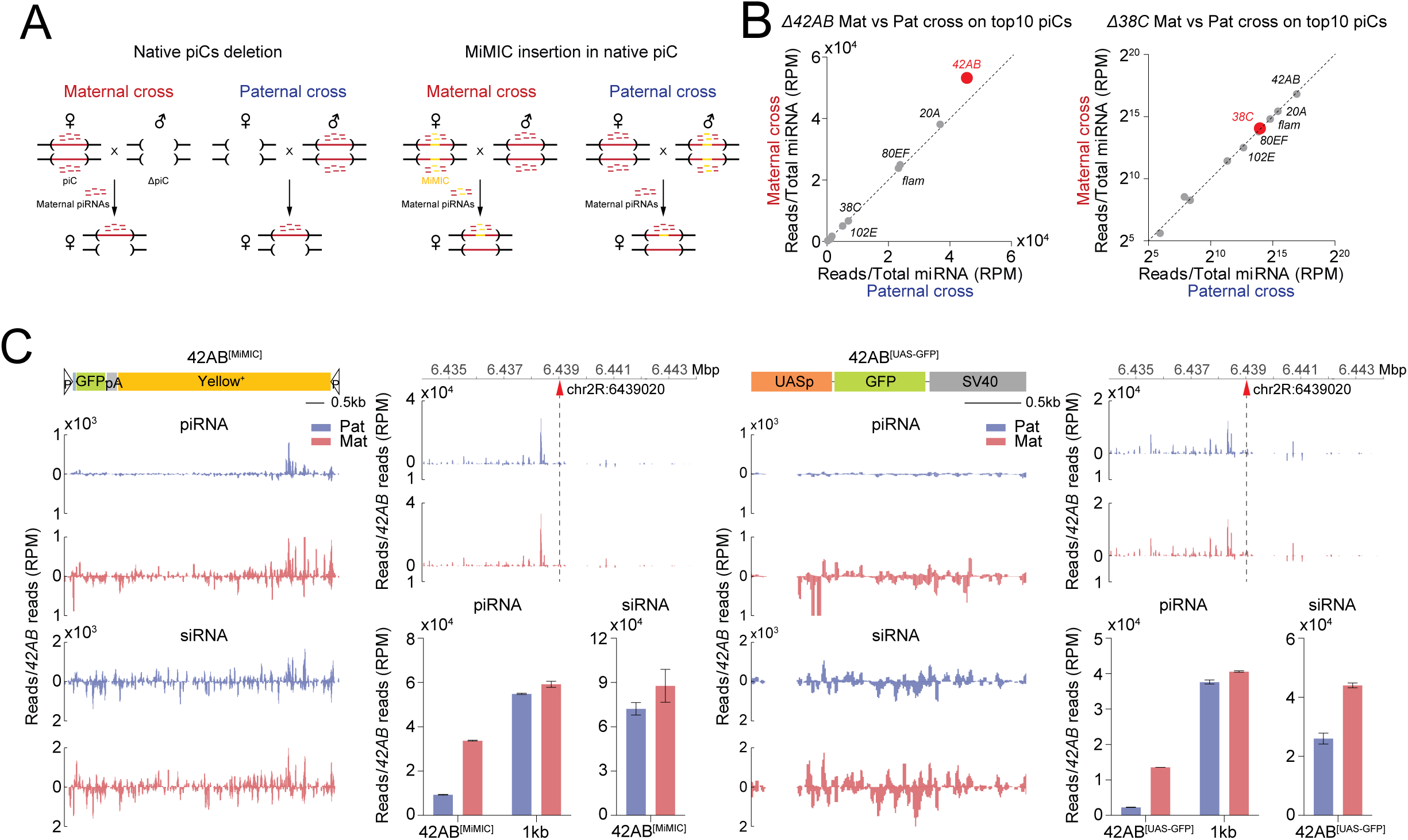
Impact of maternal piRNA on piRNA biogenesis. **A. Schematic of maternal and paternal crosses.** The native piRNA cluster is indicated by the red line within brackets, and the MiMIC insertion into the piC is shown in yellow. (Left) Flies carrying deletions of a piRNA cluster (*ΔpiC*), *42AB* or *38C*, were crossed with wild-type flies. The progeny of maternal, but not paternal cross receives cytoplasmic cluster-derived piRNAs from their mothers (Right) Flies carrying MiMIC insertion within the 42AB cluster were crossed with wild-type flies. The progeny of maternal, but not paternal cross receives MiMIC-derived piRNAs from their mothers, while both progenies inherit cluster-derived piRNAs. **B. Loss of maternally inherited piRNAs from 42AB and 38C clusters has negligible effect on piRNA levels in the progeny.** Scatter plots show piRNA abundance from the top 10 germline piRNA clusters in progenies of maternal and paternal crosses with deletion of *42AB* (left) and *38C* (right) clusters. Values represent piRNA reads uniquely mapped to each piC normalized to total miRNA reads. **C. piRNA production from cluster insertions depends on maternally inherited piRNAs.** piRNA and siRNA profiles at the 42AB^[MiMIC]^ and 42AB^[UAS-GFP]^ insertions and surrounding genomic regions in progenies of maternal (red) and paternal (blue) crosses. While the levels of piRNA derived from insertion sequences are drastically different in the progenies of maternal and paternal crosses, the levels of insertion-derived siRNA as well as piRNA from flanking sequences are similar in two progenies. Bar plots show piRNA levels in insertions and 1 kb regions flanking the insertion site in *42AB* cluster. Values were normalized to total *42AB* piRNA reads.

An alternative to using piC deletions to explore the role of cytoplasmic piRNA inheritance is employing clusters with insertion of additional sequences. Any sequence embedded in a cluster generates piRNA^17,19^. Thus, reciprocal crosses between flies that have insertions into piC and flies with non-modified clusters provide a tool to study the function of maternal piRNA generated from the inserted sequence (Fig. 1A, right panel). We took advantage of the MiMIC project that generated an insertion of a heterologous sequence in the *42AB* cluster (42AB^[MiMIC]^ allele) and employed recombination to replace the MiMIC sequence with the UASp-GFP sequence, creating the 42AB^[UAS-GFP]^ allele. Analysis of short and long RNA expression as well as chromatin structure showed that heterologous sequences (cassettes) embedded in the *42AB* cluster became its integral part and was transcribed by the non-canonical Rhi-dependent transcriptional machinery to generate piRNAs and siRNAs derived from both genomic strands (Fig. S1).

The analysis of piRNA profiles in ovaries of adult progenies of reciprocal crosses between flies with and without insertions revealed that the levels of cassette piRNAs were higher in progeny of maternal cross (i.e., where the cassette was inherited from the mother). The difference compared to the reciprocal cross was 3.6-fold (n=2, p=0.000059, two-tailed Student’s t test) and 6-fold (n=2, p=0.000021) for the MiMIC and UASp-GFP cassettes, respectively (Fig. 1C). In contrast to the large difference in piRNA amounts, we only observed a minor difference in the amount of siRNA between progenies from the maternal and paternal crosses: 1.2- and 1.7-fold for the MiMIC and UASp-GFP, respectively. Thus, piRNA, but not siRNA biogenesis in the progeny depends on maternal input of cognate non-coding RNAs. Importantly, the levels of *42AB* piRNAs from non-cassette sequences, including regions flanking the cassette insertions, were similar in progenies of the reciprocal crosses (Fig. 1C).

Thus, the two approaches used to address the role of maternal piRNA on piRNA biogenesis from the *42AB* cluster produced strikingly different and paradoxical results: complete absence of maternal *42AB* piRNA had negligible effect on piRNA biogenesis from this cluster in the progeny, while the absence of piRNAs from the heterologous insertion leads to a dramatic reduction in piRNA biogenesis from this insertion.

### piRNA encoded *in trans* can compensate for a deficiency of *cis*-piRNA

What could be the reason for the dramatic difference in the effect of maternal piRNAs in experiments that use cluster deletions and insertions? piCs harbor TEs and other repeats, which are present elsewhere in the genome. If genomic regions outside of piCs could generate piRNAs that target it, such *trans*-acting piRNAs might be able to compensate for the lack of cluster-derived *cis*-piRNA. In contrast, heterologous sequences such as the MiMIC and UASp-GFP inserted into *42AB* are targeted exclusively by *cis*-piRNA.

To explore if piCs are targeted by *trans*-acting piRNA produced from other genomic loci, we profiled small RNAs in flies with deletions of the *42AB* and *38C* clusters. Flies with *42AB* deletion retained 47.2% of piRNA reads (56.8% of distinct sequences) that mapped to the deleted locus with no mismatches (Fig. 2A). Similarly, 28.4% of piRNA reads (54.7% of distinct sequences) with perfect match to *38C* remained in flies with deletion of this cluster (Fig. 2A). Importantly, piRNAs do not have to be completely complementary to target a sequence for effective recognition as piRNAs can repress targets that have ∼ 90% sequence identity^25,26^, which for typical piRNA length of 23-26nt corresponds to 2-3 mismatches between the piRNA and its target. The analysis of piRNAs that can target the deleted sequence of the *42AB* and *38C* clusters with up to 3 mismatches yielded additional piRNAs, many of which remained present in flies with deletion of the corresponding cluster (Fig. 2A). Comparative analysis of piRNA profiles in flies with and without deletions of the *42AB* and *38C* clusters enabled us to determine the levels of *trans*-piRNAs that were produced elsewhere in the genome but might target these clusters. Importantly, *trans*-piRNAs compose 60.8% of total piRNAs targeting *42AB,* indicating that the impact of *trans*-piRNAs exceeds that of *cis*-piRNAs. *Trans*-piRNAs constitute a smaller, but still large fraction of piRNAs (34.8%) targeting *38C*.

**Figure 2.**
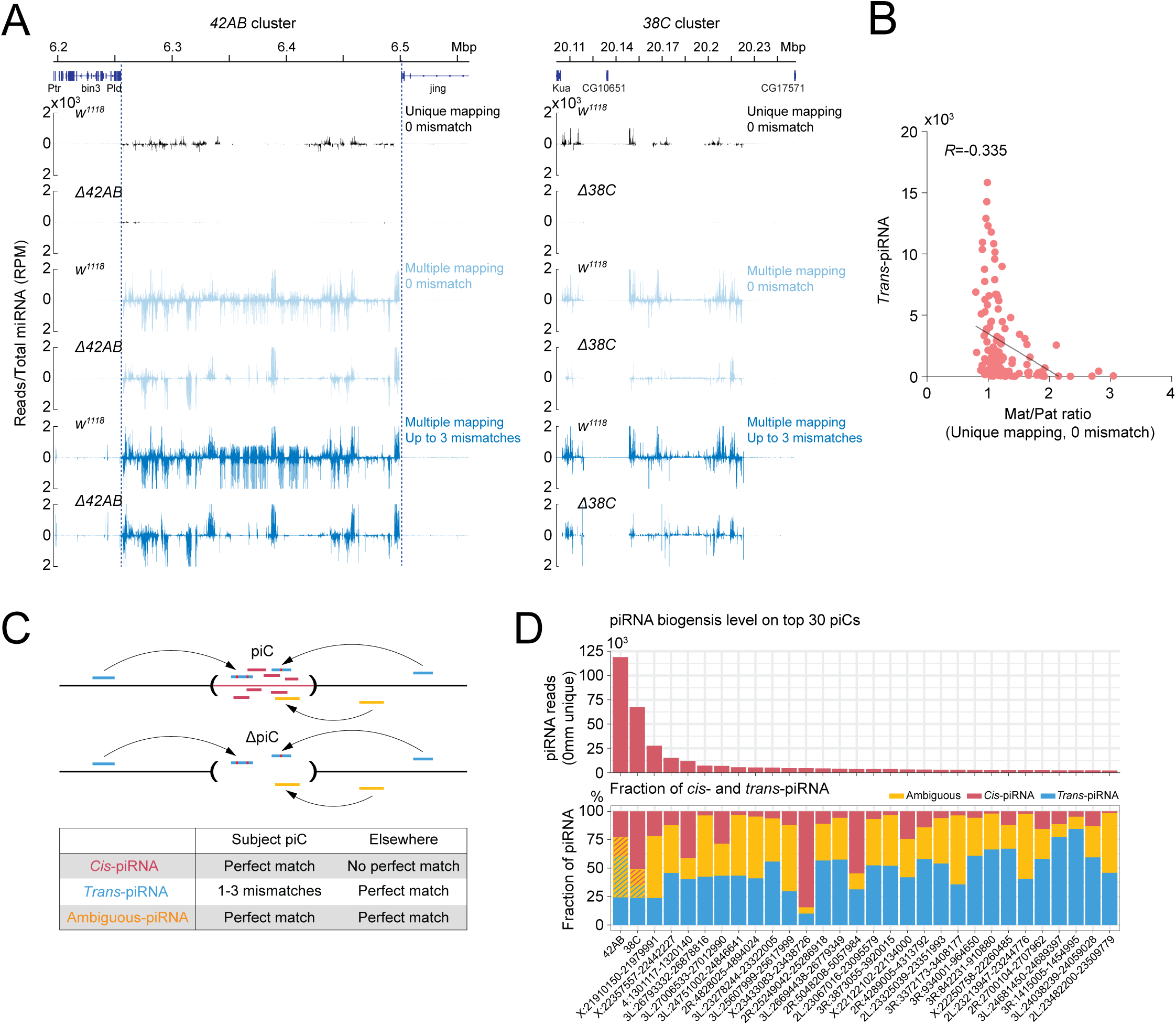
*trans*-acting piRNA compensate for a deficiency of *cis*-piRNA. **A. Abundant piRNAs are mapped to *42AB* or *38C* clusters in the flies lacking *42AB* or *38C* cluster.** Shown are piRNA profiles on *42AB* or *38C* piCs in the wild-type flies (w^1118^) and flies homozygous for deletions of these clusters. The piRNA reads were mapped either with no mismatches (0 mismatch) or allowing up to 3 mismatches. No mismatch reads are mapped to genome at unique positions (Unique mapping) or several genomic regions (Multiple mapping). The results were normalized to total miRNA in two biological replicates. **B. Correlation between abundance of *trans*-acting piRNA and sensitivity of piRNA biogenesis to impact of maternal *cis*-piRNA in *42AB* cluster.** The scatter plot show the correlation between levels of *trans*-acting piRNA targeting 1 kb genomic windows in *42AB* cluster and sensitivity of piRNA biogenesis to supply of cytoplasmically-inherited maternal cis-piRNA. The sensitivity is measured as Mat/Pat ratio, a ratio of piRNAs generated from each genomic window in two progenies, the progeny that inherited maternal cis-piRNA for *42AB* cluster and the progeny that lack *42AB* cis-piRNA (progenies of maternal and paternal crosses shown on Fig 1A, left). The levels of *trans*-acting piRNAs that recognize genomic window in *42AB* cluster with up to 3 mismatches were determined in flies that lack cis-piRNA due to deletion of *42AB*. Only genomic windows that produce more than 50 RPM of uniquely-mapping piRNA reads in paternal cross were included in the analysis. Pearson correlation coefficient (R) is indicated. **C. Categorization of *cis*- and *trans*-acting piRNA.** Dual-strand piRNA clusters can be recognized and targeted by *cis-*piRNAs (red) derived from the same cluster and *trans-*acting piRNAs (blue) encoded elsewhere in the genome. In the absence of the data obtained in flies with deletion of corresponding piC, some piRNAs cannot be unequivocally classified as *cis*- or *trans-*piRNA (yellow). In flies with deletion of piC (delta piC, bottom) *cis-*piRNA are eliminated allowing for precise determination of *trans-*piRNA. For categorization purposes in the absence of piC deletion data (table), we consider *cis*-piRNA as reads that are uniquely mapped to subject cluster with no mismatches. *Trans*-acting piRNA are reads that mapped to subject cluster with 1 to 3 mismatches and mapped elsewhere in the genome with no mismatches. Ambiguous piRNA are reads that mapped with no mismatches in the subject cluster and in addition elsewhere in the genome. **D. Abundance of *cis*- and *trans*-piRNAs for the top piRNA clusters.** The upper plot shows amounts of *cis*-piRNA produced by the top 30 piRNA clusters in the ovaries of wild-type flies. The bottom plot shows for the same clusters proportions of *cis*-, *trans*-, and ambiguous piRNA as defined on Fig. 2C. The two biological replicates were averaged. For two clusters, *42AB* and *38C*, where deletion mutants lacking cis-piRNA are available we determined the fractions of ambiguous piRNAs which are derived from genuine cis-and trans-piRNA (red and blue stripes, respectively).

Analysis of *trans*-piRNAs revealed that they are unequally distributed along the *42AB* and *38C* clusters, with some regions having a large density of *trans*-piRNAs and others lacking them (Fig. 2A). If *trans*-piRNAs compensate for the lack of maternal *cis*-piRNAs, then regions that lack *trans*-piRNAs should be affected more severely in the progeny of paternal cross. To explore this hypothesis, we analyzed piRNA biogenesis in the progenies of paternal and maternal crosses in genomic windows along the *42AB* cluster. Though the overall difference in *42AB*-derived piRNA between paternal and maternal crosses was small (14%) (Fig. 1B), a few genomic windows show much stronger, up to 3-fold, loss of piRNAs in the progeny from the paternal cross. Remarkably, all these windows had no or very little *trans*-piRNAs that could target them (Fig. 2B). Overall, we observe a negative correlation between the level of *trans*-piRNAs targeting any given genomic window and the maternal-to-paternal piRNA ratio in the progeny (R=-0.335), suggesting that uneven *trans*-piRNAs profiles along clusters lead to different piRNA biogenesis efficiency.

Investigation of piRNA profiles in flies with deletion of the *42AB* and *38C* clusters revealed that these clusters are targeted by *trans*-piRNAs derived from other genomic regions. However, this approach cannot be generalized to other piCs for which deletions are not available. Hence, we sought to develop a general approach that would allow us to estimate the levels of *cis* and *trans*-piRNAs for any cluster (Fig. 2C). The minimum conservative estimate of *cis*-piRNA generated by a subject cluster is the number of piRNAs that map to this cluster and nowhere else in the genome (perfect unique mappers). The minimum estimate of *trans*-piRNAs that target a subject cluster is the pool of piRNAs that have 1 to 3 mismatches to the sequence of subject cluster, while having perfect mapping elsewhere, indicating that they are transcribed from other genomic regions. In addition to these two groups, piCs are targeted by a pool of piRNAs with ambiguous origin: such piRNAs have perfect match to a subject cluster sequence and they also map perfectly elsewhere in the genome, making it impossible to unambiguously determine their origin (Fig. 2C). We calculated these three fractions for the top 30 dual-strand piRNA clusters (Fig. 2D). The analysis revealed that all 30 piCs have large fractions of *trans*- and ambiguous piRNAs. For different piCs the fraction of *trans*-piRNAs varies from 10% to 84.6% with a mean value of 47.5% and standard deviation of 16.5%. The fraction of ambiguous piRNAs varies from 5.42% to 60.6% with a mean value 36.1% and a standard deviation of 15.8%. Remarkably, for 14 out of the top 30 clusters, the minimum estimate of *trans*-piRNAs exceeded that of the combined amounts of *cis* and ambiguous fractions. For 3 clusters, the minimum estimate of *trans*-piRNA exceeded that of *cis* and ambiguous fractions by more than 2-fold. Notably, the majority of piCs the fractions of *trans*-piRNAs is higher than for *42AB* and *38C*. Overall, our analysis indicates that dual-strand clusters are targeted by abundant *trans-*piRNAs that are produced from other genomic loci.

### Extensive crosstalk between piRNA clusters promotes piRNA biogenesis

Where do *trans*-piRNAs targeting piCs come from? To understand the origin of *trans*-piRNAs, we analyzed piRNAs targeting each cluster while being produced from other clusters. For each pair of clusters, we determined the number of piRNAs produced by the donor cluster (with 0 mismatches) that target the recipient cluster (with one to three mismatches) (Fig. 2C). The result of this analysis is a matrix with columns corresponding to clusters generating the piRNAs, and rows of clusters targeted by these piRNAs. In addition to visualizing the normalized read number (RPM) of *trans*-piRNAs (Fig. 3A, left), we performed independent scaling for each recipient cluster to determine the fractions of *trans*-piRNAs from donor clusters (Fig. 3A, right). The analysis revealed extensive crosstalk between individual piCs with almost all clusters participating in interactions as both donors and recipients of *trans*-piRNAs. Furthermore, the vast majority of clusters are targeted by *trans*-piRNAs generated from multiple other clusters. As a result of such extensive interactions, the relative contribution of each individual donor cluster to the pool of *trans*-piRNAs targeting a recipient cluster is typically small (with notable exceptions of large contributions of *42AB* and a few other clusters described below). Indeed, in 80.4% of all pairwise interactions, individual donor clusters contribute less than 5% of the total *trans-*piRNAs received by an acceptor cluster. Thus, our global analysis indicates profound interconnectivity of piCs, with clusters forming one common network with direct links between individual clusters.

**Figure 3.**
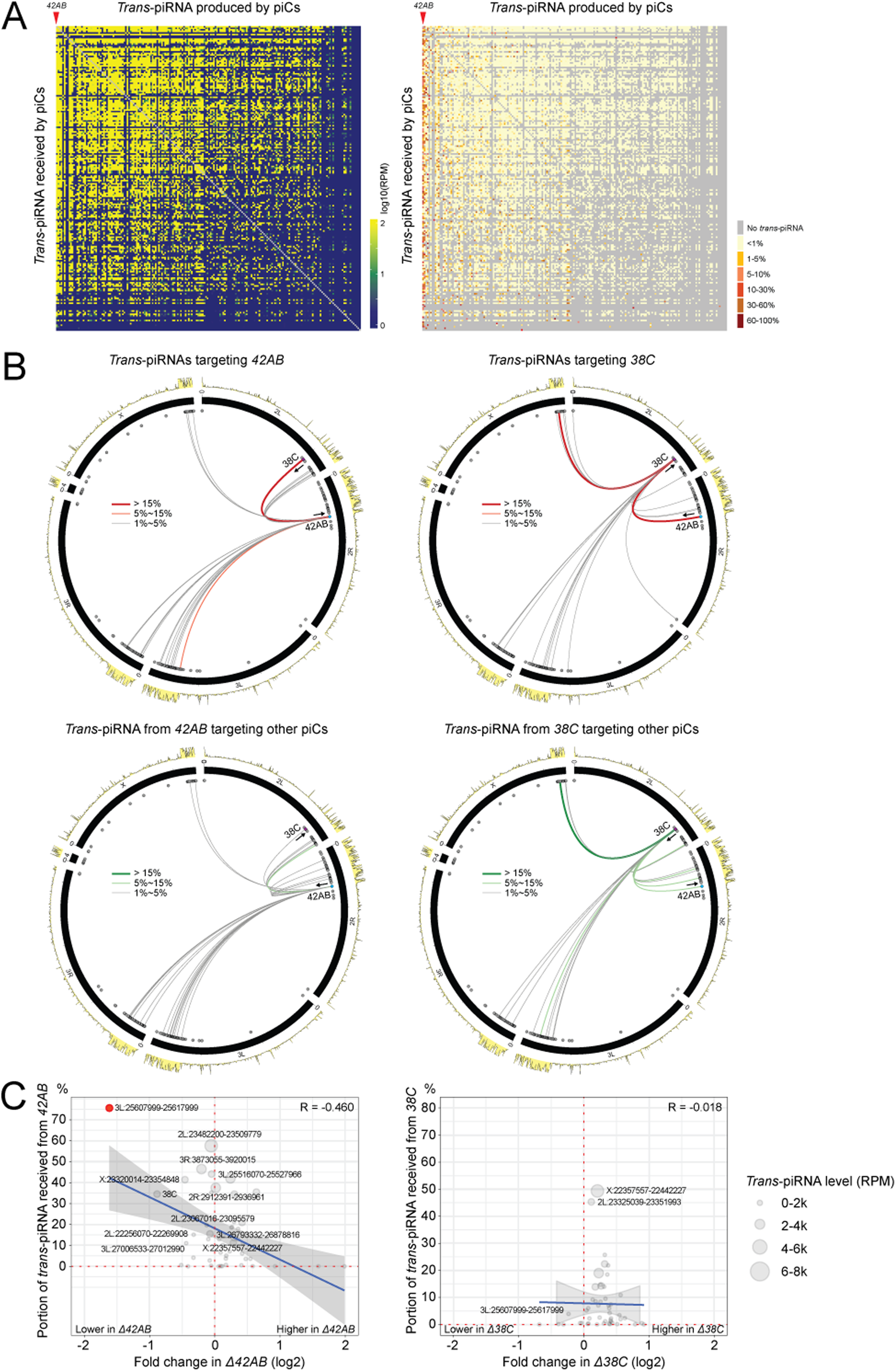
Extensive crosstalk between piRNA clusters promotes piRNA biogenesis. **A. *Trans*-acting piRNAs connect individual piCs into integrated network.** The symmetric correlation matrices show interaction of individual clusters through *trans*-piRNA with piCs as rows and columns. Columns represent piCs generating *trans-*piRNAs, and rows represent recipient clusters targeted by these piRNAs. The left matrix shows the levels of *trans-*piRNA that connect each pair of clusters as read counts (RPM). The right matrix adds a scaling of *trans-*piRNA levels by the total *trans*-piRNA targeting each recipient cluster resulting in a matrix that shows proportional contribution of *trans-*piRNA from different clusters to each recipient cluster. **B. *Trans*-piRNA interactions involving the *42AB* and *38C* clusters.** Circos plots show *D. melanogaster* chromosomes as black ranges. The outer yellow track shows genomic origin of uniquely mapped piRNAs expressed in wild-type flies. piCs are marked by grey dots, except for clusters *42AB* and *38C* in blue and purple, respectively. Lines linking clusters represent interactions between clusters through *trans*-piRNAs. On the top plots the color of the lines shows the percentage of *trans*-piRNA generated by different clusters and targeting *42AB* and *38C*. On the bottom plots, the color of the lines shows *trans*-piRNAs produced by *42AB* and *38C* and targeting other clusters. Grey lines: contribution >= 1% and < 5%, thin red/green: contribution >= 5% and < 15%, thick red/green: contribution >15%. Arrow direction towards cluster: reception of *trans*-piRNA, arrow direction opposed to cluster: generation of *trans*-piRNA by the cluster. **C. Loss of *trans*-acting piRNAs impairs piRNA biogenesis at recipient clusters.** Scatter plot shows the effect of deletions of *42AB* (left) and *38C* (right) clusters on piRNA biogenesis from other clusters (x-axis). The y-axis shows the proportion of total *trans*-piRNAs targeting recipient clusters produced by *42AB* (left) and *38C* (right). The size of each point indicates the total levels of *trans*-piRNA received from *42AB* and *38C*. The only cluster that exhibited more than a two-fold reduction due to the 42AB deletion is highlighted in red. Pearson correlation coefficient (R) is indicated.

The pairwise matrix revealed that the most active clusters such as *42AB* and *38C* are prolific producers of *trans*-piRNAs targeting other clusters. Indeed, 42 different clusters receive more than 30% of their total *trans*-piRNAs from *42AB*, while *38C* contributes more than 30% of *trans*-piRNAs for 14 piCs. Remarkably, *42AB* and *38C* generate *trans*-piRNAs targeting each other (Fig. 3B). *42AB* is the top provider of *trans*-piRNAs to *38C*, contributing 34.5% of all *trans*-piRNAs targeting *38C.* However, these piRNAs represent only 1.56% of the total *trans*-piRNAs produced by *42AB.* Reciprocally, *38C* is the top *trans*-piRNA provider to *42AB*, contributing 19% of trans-piRNAs targeting this cluster. From the *38C* perspective, *42AB*-targeting piRNAs compose 8.3% of the total *trans*-piRNA pool produced by *38C.* Thus, though the majority of individual pairwise interactions between clusters are weak, *42AB* and to some extent *38C* are distinct as they provide large fractions of *trans*-piRNAs to many recipient clusters, including to each other (Fig. 3B).

Our analysis revealed an extensive network of cluster interactions dominated by small cumulative contributions of individual donor clusters, suggesting that deletion of any single cluster might have a negligible impact on the expression of its target clusters. The possible exception is *42AB,* which contributes a significant fraction of *trans*-piRNAs targeting several other clusters (Fig. 3A and 3B). To explore the functional effect of *42AB* on other clusters, we analyzed piRNA expression in *42AB* deletion flies and compared it with that of *38C* deletion. Remarkably, 17 of the top 30 clusters (56%) had lower expression in flies with *42AB* deletion, while only 6 had lower expression upon *38C* deletion (Fig. 3C). For most affected clusters, the effect was relatively weak (less than 2-fold). The prominent exception is cluster *chr3L:25,607,999-25,617,999,* which had ∼3-fold lower expression upon *42AB* deletion. Remarkably, this cluster also had the highest fraction (75.6%) of recipient *trans*-piRNAs produced by *42AB*. *38C* – for which 34.5% of *trans*-piRNAs are produced by *42AB* – showed 1.8-fold (−0.8 log2 fold change) reduction of expression in the *42AB* deletion mutant. Notably, the reduction in expression of these two clusters is much stronger than the reduction in expression of *42AB* itself in the progeny of the paternal cross (Fig. 1B). This analysis revealed a correlation between the reduction of clusters expression upon *42AB* deletion and the fraction of *42AB*-derived *trans*-piRNA targeting these clusters (*Pearson’s* correlation: *R* = −0.46). Thus, the lack of piRNAs from prolific clusters such as *42AB* can have a significant impact on the expression of other clusters. Overall, the analysis of cluster interactions through *trans*-piRNAs and studies of the effects of *42AB* deletion on expression of other clusters demonstrate that dual-strand piCs are involved in an extensive network of mutually supporting interactions that might be described by the ‘one for all and all for one’ motto.

### Maternal piRNAs are dispensable for transcription of piRNA precursors but essential for their cytoplasmic processing

What is the mechanism by which maternal piRNAs activate piRNA biogenesis in the progeny? Previous work using transgenic clusters implicated maternal piRNAs in promotion of non-canonical transcription (NCT) of piRNA precursors through deposition of the H3K9me3 mark, followed by binding of Rhi, which forms a complex that recruits other proteins required for NCT^10,16,17^. *42AB* cluster is enriched in H3K9me3 and Rhi and piRNA production strongly depends on Rhi and its protein partners implicated in NCT^7,8,14^. Heterologous sequences (MiMIC and UASp-GFP) embedded in *42AB* have the same properties (Fig. S1), indicating that Rhi-dependent NCT is responsible for their expression and piRNA biogenesis.

To explore the role of maternal piRNA in Rhi-dependent NCT, we analyzed chromatin over and nascent transcripts from the inserted casette in progenies of maternal and paternal crosses. Nascent transcripts from two genomic strands are detected in germ cell nuclei in both progenies. In contrast, no cytoplasmic RNA from either strand was detected in either progeny (Fig. 4A). Furthermore, quantitative analysis of FISH signals revealed no difference between nascent cassette RNA levels in the progenies of the reciprocal crosses (Fig. 4B). Similarly, ChIP-qPCR showed no significant difference in H3K9me3 (Pat=0.92, Mat=0.99, p=0.06) and Rhi occupancy (Pat=1.03, Mat=1.07, p=0.36) (Fig. 4C). The combined analysis of nascent transcripts and chromatin indicate that maternal piRNAs do not affect deposition of Rhi and cassette NCT. Thus, while Rhi-dependent NCT is essential for piRNA biogenesis from cassettes inserted into *42AB* (Fig. S2A), this process is not affected by cytoplasmic piRNA inheritance, indicating that maternal piRNAs promote piRNA biogenesis by a different mechanism.

**Figure 4.**
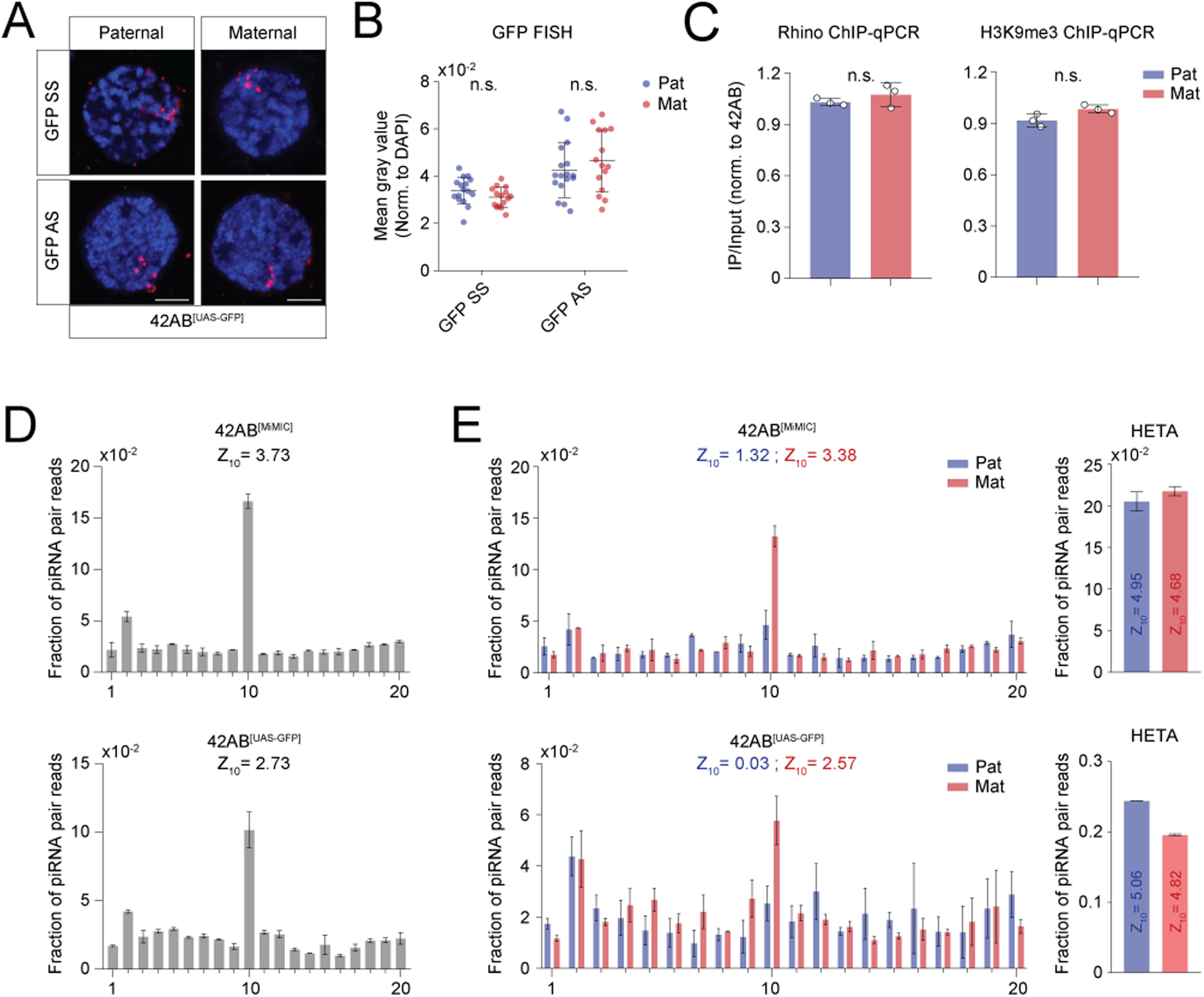
Maternal piRNAs are dispensable for transcription of piRNA precursors from heterologous insertions into *42AB* cluster but essential for their cytoplasmic processing. **A. Nascent transcripts from UASp-GFP insertion in *42AB* cluster are detected in nuclei of germline cells in maternal and paternal crosses.** RNA FISH was performed separately using probes targeting GFP sense and antisense transcripts (red) in progenies of maternal and paternal crosses shown on Fig 1A, right. Scale bar represents 5μm. Statistical significance is estimated by two-tailed student’s t-test; n.s.> 0.05. **B. Quantitative analysis shows no significant difference in UASp-GFP nascent transcript levels between progenies of maternal and paternal crosses.** Dot plots shows intensities of GFP sense and antisense RNA FISH signal in individual nuclei. The number of nuclei is approximately 15 for each experiment. FISH images were taken using the same settings. **C. The levels of Rhi and H3K9me3 on UASp-GFP insertion are similar in progenies of maternal and paternal crosses.** Rhi and H3K9me3 ChIP-qPCR were performed on ovaries from the progenies of maternal and paternal cross. Results were measured in three biological replicates and normalized to *42AB* region (chr2R: 6,323,128 – 6,323,219). Error bars indicate standard deviation of three biological replicates. Statistical significance is estimated by two-tailed student’s t-test; n.s.> 0.05. **D. Heterologous insertions in *42AB* cluster exhibit a strong signature of ping-pong amplification.** The analysis of the ping-pong signature was performed on MiMIC and UASp-GFP insertions. **E. The ping-pong amplification depends on cytoplasmic inheritance of maternal piRNAs.** The analysis of the ping-pong signature was performed on MiMIC and UASp-GFP insertions in progenies of maternal and paternal crosses. As a control, the ping-pong signature was analyzed on HETA transposable element.

Next, we explored if maternal piRNAs affect the cytoplasmic processing of piRNA precursors into mature piRNAs. In the cytoplasm, piRNAs are amplified through two linked mechanisms, the phasing pathway and the ping-pong cycle^2,27,28^. We did not detect phasing pattern on MiMIC and UASp-GFP sequences in progenies of the maternal and paternal crosses (Fig. S2B). In contrast, piRNAs produced by the MiMIC and UASp-GFP cassettes revealed strong signature of ping-pong processing, enrichment in characteristic pairs of complementary piRNAs with 5’ends separated by 10nt (Fig: 4D). The ping-pong signature was also prominent in progenies of the maternal cross (Fig. 4E). In fact, the ping-pong scores (Z_10_) were similar in the progeny of the maternal crosses and their mothers, 3.38 and 3.73 (MiMIC), 2.57 and 2.73 (UASp-GFP), respectively. In contrast, no ping-pong signature was detected in the progeny of the paternal cross, indicating the collapse of the ping-pong cycle in the absence of maternal piRNAs. Taken together, our analysis of piRNA biogenesis suggests that cognate maternal piRNAs are dispensable for transcription of piRNA precursors but strongly promote their ping-pong processing in the cytoplasm.

### Canonical transcription inside piCs interferes with non-canonical transcription

Insertion of heterologous sequences in piCs models the process of TE insertions^23,29,30^. However, unlike promoterless sequences used in previous experiments, natural transposons carry regulatory elements that drive their own transcription. To test the trap model of piRNA pathway adaptation, we studied the effect of inserting a transcriptionally active gene into a piC. The UASp-GFP cassette carries a UASp promoter that can be activated by the yeast GAL4-VP16 protein. We explored expression of UASp-GFP inserted in *42AB* cluster upon GAL4 induction in progenies of the two reciprocal crosses that are genetically identical, but distinct in their inheritance of maternal piRNAs (Fig. 5A). Upon expression of GAL4 under the control of the maternal α-Tub67C promoter (MTG), GAL4 activates RNA pol II transcription, leading to expression of GFP protein in germ cells (Fig. 5B). The timing of MTG-GAL4 induced transcription and NCT of *42AB* cluster overlaps, as both are detected in germ cells starting in stage 2 to 3 egg chambers and continues throughout oogenesis^31,32^.

**Figure 5.**
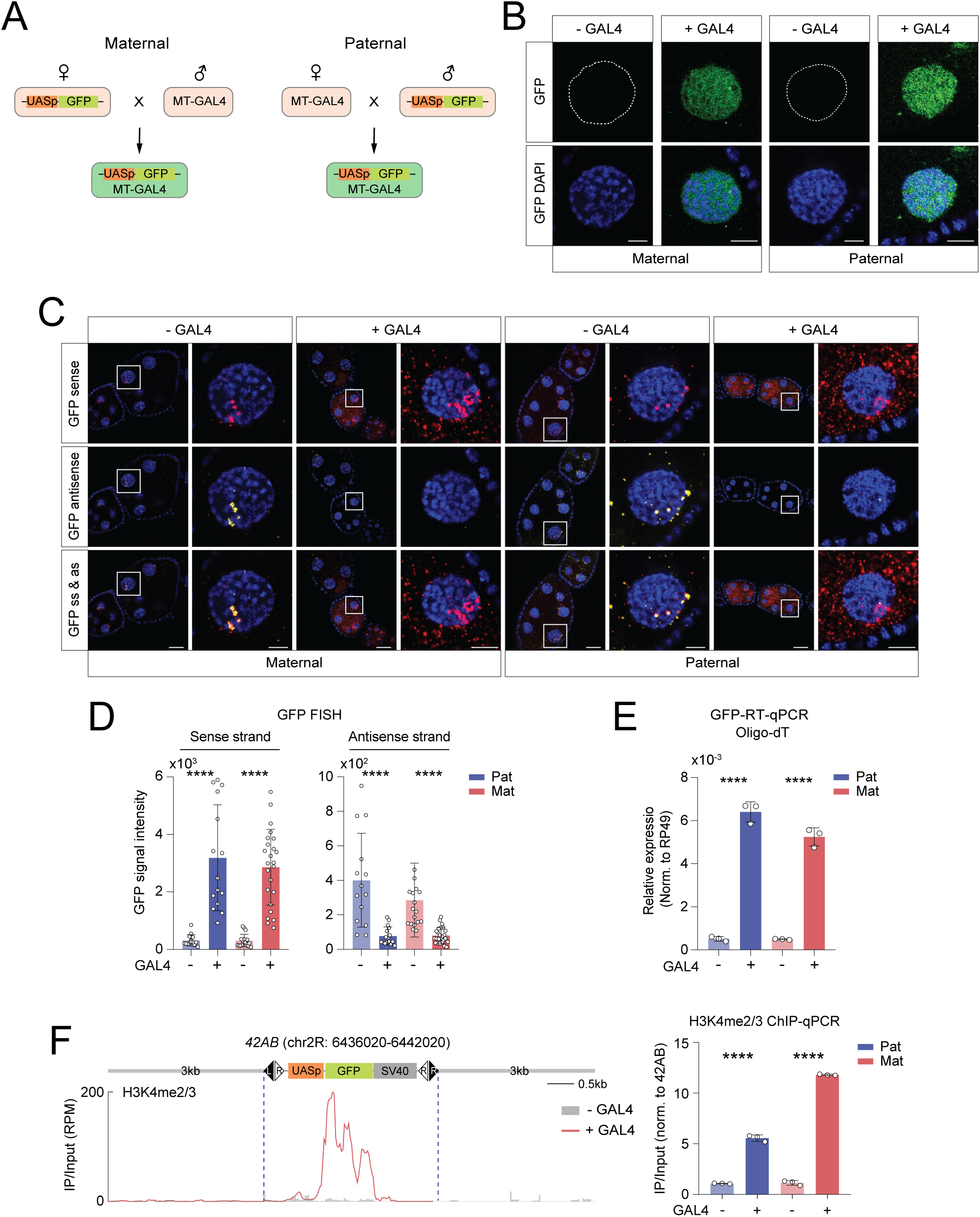
The canonical transcription inside *42AB* cluster interferes with non-canonical transcription of the piRNA precursors. **A. The scheme of maternal and paternal crosses between flies with UASp-GFP insertion in *42AB* cluster and MT-GAL4 driver.** **B. UASp-GFP insertion inside *42AB* cluster is activated by GAL4 induction and produces functional protein.** GFP fluorescence was measured with and without MT-GAL4 induction in progenies of maternal and paternal crosses. GAL4-induced transcription leads to GFP expression in progenies of both crosses. Scale bar represents 5μm. **C. GAL4-induced transcription dominates and represses NCT.** FISH for simultaneous detection of sense (red) and antisense (yellow) GFP RNA shows that activation of UASp promoter by GAL4 triggers accumulation of sense GFP RNA in cytoplasm and simultaneous loss of antisense RNA indicating collapse of NCT. The similar results were obtained in progenies of maternal and paternal crosses. Scale bar is 20 μm and 5 μm for egg chamber and single nurse cell nuclei, respectively. **D. Quantification of nascent RNAs in individual nuclei shows suppression of NCT by GAL4-induced transcription.** Bar graphs show the quantification of GFP sense and antisense FISH signal intensity in individual nurse cell nuclei. GAL4 induction increases the levels of sense RNA and decreases the levels of antisense RNA in paternal and maternal crosses. The number of nuclei is at least 15 in each experiment. Images were taken using the same settings. Statistical significance is estimated by two-tailed student’s t-test; **** p<0.0001. **E. The polyadenylated GFP RNA accumulated upon GAL4 induction.** Polyadenylated GFP RNA was measured by RT-qPCR using oligo dT primer for reverse transcription. *rp49* mRNA levels were used for normalization. Error bars indicate the standard deviation of three biological replicates. Statistical significance is estimated by two-tailed student’s t-test; **** p<0.0001. **F. Active chromatin H3K4me2/3 mark accumulates on the UASp-GFP insertion upon GAL4 induction.** (Left) The ChIP-seq profiling shows no H3K4me2/3 mark in the absence of GAL4 induction, while high levels of the mark appear at the 5’ end of the UASp-GFP insertion upon induction. The mean values from two biological replicates are shown. (Right) H3K4me2/3 ChIP-qPCR experiments were performed in three biological replicates and normalized to rp49 region. Error bars indicate standard deviation of three biological replicates. Statistical significance is estimated by two-tailed student’s t-test; **** p<0.0001.

We further explored cassette expression upon GAL4 induction by analyzing nascent transcripts and cytoplasmic mRNA as well as chromatin marks. Without GAL4 induction, transcripts from the two genomic strands are co-localized and present almost exclusively in the nucleus. Upon induction, sense-strand RNA increased in the nucleus (10-fold and 11.9-fold in progenies of the maternal and paternal crosses, respectively) and was abundant in the cytoplasm (Fig. 5C and 5D). This pattern was identical to GAL4-induced expression of the cassette inserted in a non-cluster locus and typical for mRNA expression. At the same time, induction led to a decrease in antisense nascent RNA levels (3.5-fold and 5.1-fold in progenies of maternal and paternal crosses, respectively) (Fig. 5C and 5D). Interestingly, we observed heterogeneity in response to GAL4 induction even between individual cells within a single egg chamber. Upon induction, a subset of cells displayed strong cytoplasmic sense GFP signal and almost undetectable levels of antisense RNA. Other cells retain a similar pattern of expression to that observed without GAL4 induction: low levels of cytoplasmic sense RNA and high levels of nuclear antisense signal (Fig. S3A). Thus, upon GAL4 induction transcription is drastically reorganized in some, but not all cells: sense mRNA generated by CT appears in the cytoplasm, while antisense RNA decreases indicating suppression of NCT. Importantly, changes caused by activation of CT were independent of the cytoplasmic inheritance of maternal piRNAs, as they occurred in progenies of both of the reciprocal crosses.

Expression of GAL4 triggered canonical transcription inside *42AB* cluster that produced GFP mRNA. First, GFP protein is detected in the germline starting at stage 2, indicating that sense mRNA is translated (Fig. 5B). Second, RT-qPCR detected strong increase in the levels of polyadenylated cassette RNA: 10.6-fold and 12.5-fold in progenies of maternal and paternal crosses, respectively (Fig. 5E). As piRNA precursors produced by NCT lack polyA-tail^8,14^, the increase in polyadenylated RNA indicates a switch from non-canonical to canonical transcription. Finally, upon GAL4 induction, we detected strong accumulation of H3K4me2/3, the chromatin mark associated with initiation of CT, on the cassette’s promoter sequence (Fig. 4F). Taken together, these results indicate that in a subset of cells, activation of CT led to expression of mRNAs and collapse of NCT that produced piRNA precursors.

### Canonical transcription removes Rhi and reduces piRNA production

NCT of piRNA precursors depends on a complex of proteins that is anchored on chromatin by Rhi bound to the H3K9me3 mark^8,10,11^. We explored whether CT of the UASp-GFP cassette embedded in the *42AB* cluster triggered changes in H3K9me3 and Rhi occupancy. Independent ChIP-seq and ChIP-qPCR experiments indicated that GAL4-induced transcription had no significant effect on the level of H3K9me3 on the cassette and flanking sequences (Fig. 6A). However, induction of CT caused 2- to 3.4-fold decrease in the levels of Rhi over the GFP sequence in both reciprocal crosses (Fig. 6A). Furthermore a 1.6- to 1.8-fold reduction of Rhi occupancy was also observed over 1 kb cluster sequences flanking the GFP insertion. Thus, the collapse of NCT caused by CT correlates with a decrease in Rhi occupancy. Considering the essential role of Rhi in recruiting the NCT machinery^7–9,12,14^, removal of Rhi by CT might be a direct cause of NCT collapse.

**Figure 6.**
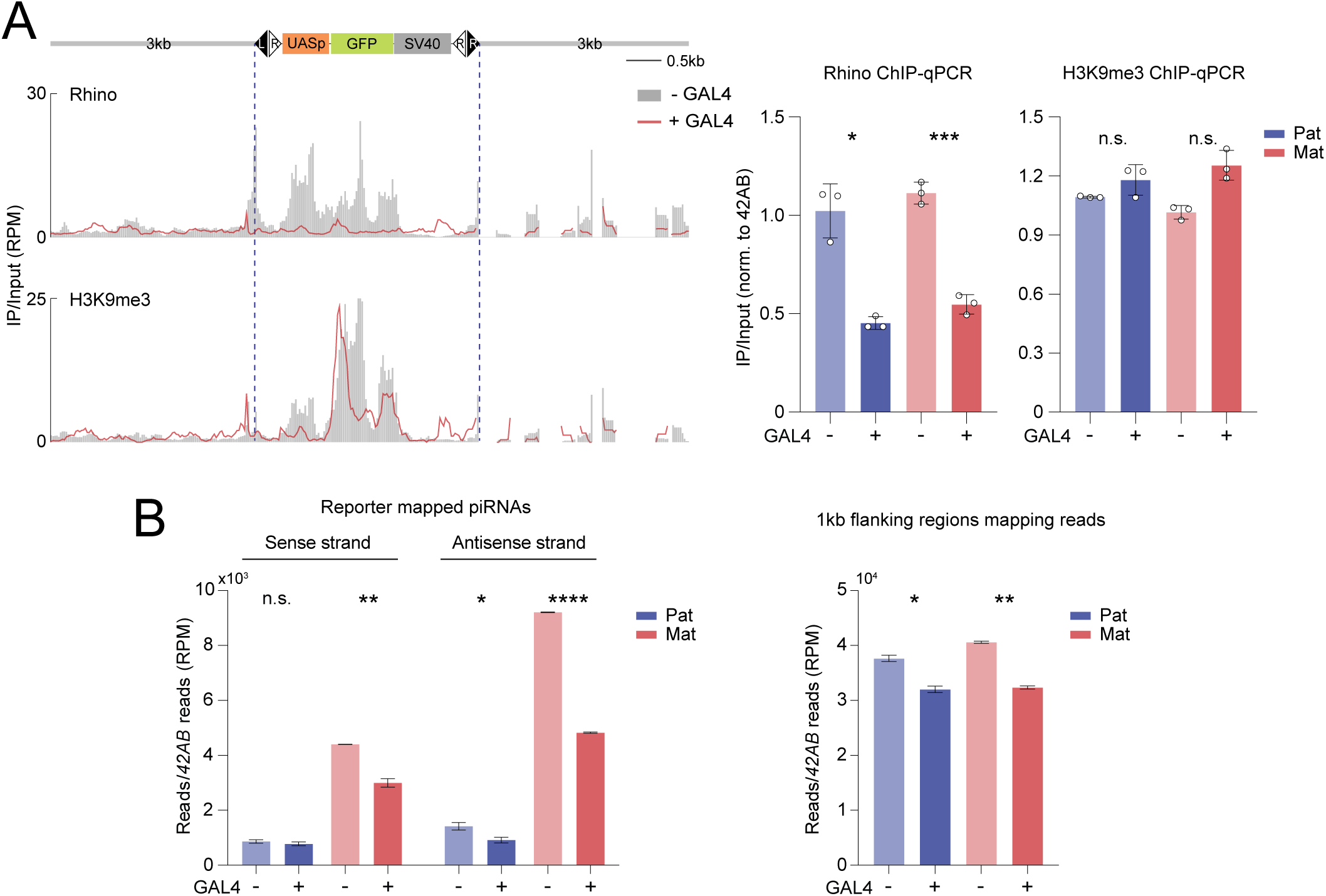
Canonical transcription in *42AB* cluster removes Rhi and reduces piRNA production. **A. GAL4-induced transcription in *42AB* removes Rhi without affecting the levels of H3K9me3.** (Left) The ChIP-seq profiles of Rhi and H3K9me3 over UASp-GFP insertion and 3 kb flanking regions in maternal cross. The profiles are the mean of two biological replicates. (Right) ChIP-qPCR for Rhi and H3K9me3 occupancy on UASp-GFP cassette were performed in three biological replicates and normalized to *42AB* region (chr2R: 6,323,128-6,323,219). Error bars indicate standard deviation of three biological replicates. Statistical significance is estimated by two-tailed student’s t-test; n.s.> 0.05, * p<0.05, *** p<0.001. **B. piRNAs derived from the UASp-GFP are decreased upon GAL4 induction.** The bar graph shows piRNAs corresponding to sense and antisense strand of UASp-GFP and 1 kb flanking sequences in maternal (red) and paternal (blue) crosses. The results were normalized to *42AB* piRNA reads. Error bars indicate the standard deviation of two biological replicates. Statistical significance is estimated by two-tailed student’s t-test; n.s.> 0.05, * p<0.05, ** p<0.01, **** p<0.0001.

Analysis of piRNA profiles revealed that induction of CT causes a decrease of piRNA biogenesis: cassette piRNA levels were reduced upon GAL4 induction 1.7-fold (p=0.0005) and 1.3-fold (p=0.04) in the maternal and paternal crosses, respectively. piRNAs corresponding to both strands were decreased upon induction, but the effect was stronger for antisense piRNAs (Fig. 6B). piRNAs generated from flanking cluster sequences were not strongly affected, indicating that the effect does not efficiently spread outside of the cassette (20% p=0.001 and 15% p=0.01 decrease in progenies of the maternal and paternal crosses, respectively) (Fig. 6B). Thus, piRNA profiling confirmed that CT causes local suppression of NCT that generates piRNA precursors. While maternal piRNAs do not affect nascent transcripts and chromatin structure (Fig. 5D and 6A), they had a drastic effect on the amounts of mature piRNAs. Indeed, the progeny of the maternal cross expressed 6-fold (p=0.00002) more cassette piRNA than the progeny of the paternal cross (Fig: 6B). Signature of ping-pong processing was observed between piRNAs mapping to the cassette in the progeny of the maternal (Z_10_ = 3.35) but not the paternal (Z_10_ = 1.17) cross. Thus, piRNA profiling revealed that when the insertion is transcriptionally active cytoplasmic inheritance of maternal piRNAs promotes ping-pong processing, causing a large increase in mature piRNAs without changing transcription of piRNA precursors.

Overall, our results indicate that a promoter-carrying gene inserted into a piC retains its capacity to be transcribed and to produce functional mRNA and protein. Furthermore, canonical transcription triggers the removal of Rhi and the collapse of NCT and piRNA biogenesis, challenging the belief that TEs are easily trapped when they land in piCs.

## Discussion

### piRNAs act across time and space: *trans-*acting trans-generational piRNAs ensure robustness of piRNA biogenesis and TE repression by connecting individual piCs into a functional network

How cellular transcripts are selected and channeled into piRNA processing is a central unresolved problem. Studies of two different transgenic clusters active in the *Drosophila* germline suggested that cytoplasmically inherited piRNAs transmitted from mother to progeny are essential for maintaining piRNA biogenesis^10,16,17^. However, whether the activity of endogenous piCs also depends on trans-generational inheritance of piRNAs remained unknown. We used two complementary approaches, deletions of two endogenous clusters obtained from the Teixeira and Lehman labs^33^ and integration of unique sequences into a piC, to explore the role of maternal piRNAs in biogenesis from native dual-strand clusters. Surprisingly, the two approaches produced opposite results. piRNA biogenesis was minimally affected in the progenies of mothers that lack the *42AB* or *38C* cluster (Fig. 1B). Similar observations were recently reported by Rice et al.^34^, who found that loss of maternal piRNAs did not significantly affect piRNA production at endogenous clusters. In contrast, maternally inherited piRNAs were critical for piRNA biogenesis when the MiMIC insertion in *42AB* cluster was used (Fig. 1C). How can these as well as previous results be reconciled into a cohesive model? The MiMIC insertion can only be targeted by piRNAs produced from this locus *in cis*. In contrast, natural cluster sequences are targeted by *trans*-acting piRNAs encoded elsewhere in the genome as the repetitive composition of piCs provides a basis for extensive cross-targeting through shared TE sequences (Fig. 3A). Importantly, we found that the amount of *trans*-piRNAs exceeds the amount of *cis*-piRNAs for the majority of piCs, suggesting that *trans*-acting piRNAs have a bigger impact on piRNA biogenesis from each individual cluster than its own, *cis-*generated piRNAs. Together, these results indicate that cytoplasmically inherited maternal piRNAs are essential to activate piRNA biogenesis on piCs in the next generation, however, the lack of *cis-*piRNAs can be compensated by maternal *trans-*piRNAs derived from other genomic loci.

Our analysis revealed that dual-strand piCs form an extensive network of interactions where *trans*-acting piRNAs derived from one piC target multiple other piCs (Fig. 3A). The result of these interactions is a highly integrated system that operates under a “one for all, and all for one” principle providing robustness to the whole system and redundancy between individual piCs (Fig. 3). Indeed, as relative contribution of each individual cluster to piRNAs targeting any other cluster is small, deletion of an individual cluster has negligible effect on piRNA biogenesis from other clusters. Nevertheless, we observed a rare case when a large fraction of *trans-*acting piRNAs targeting a specific cluster is derived from one cluster, *42AB*. Importantly, in this case deletion of *42AB* leads to a strong decrease in piRNA biogenesis from the recipient cluster demonstrating that *trans*-acting piRNAs are critical for maintaining piRNA biogenesis (Fig. 3C). The robustness provided by an integrated network of piCs is not restricted to piRNA biogenesis, but likely extends to the main function of the system, repression of TEs, as loss of individual piCs can be buffered by the remaining clusters through shared sequence homology and *trans*-piRNAs. Consistent with this, deletion of major clusters such as *42AB* and *38C* was shown to have a negligible effect on TE silencing and germline development^33^, while disruption of the whole piRNA pathway has dramatic effects^2,35^.

### Maternal piRNAs are required for transcription of piRNA precursors and their cytoplasmic processing

What are the molecular mechanisms by which cytoplasmically inherited piRNAs trigger piRNA biogenesis in the next generation? Previous studies that employed transgenic piCs suggest that maternal piRNAs promote piRNA biogenesis by two distinct mechanisms^16,17^. First, maternal piRNAs ensure NCT of piRNA precursors through installation of the heterochromatic H3K9me3 mark and recruitment of a complex containing the Rhi protein that directly binds this mark. Second, maternal piRNAs were shown to enhanced cytoplasmic processing of piRNA precursors by initiating the ping-pong amplification cycle. Here, we found that maternal piRNAs that target a MiMIC insertion in the endogenous *42AB* cluster strongly promote cytoplasmic ping-pong amplification, however, they have no observable effect on transcription of precursors and on H3K9me3 and Rhi occupancy on chromatin (Fig. 4). The difference is likely explained by the nature of the target locus: in previous studies, the entire transgenic cluster was depended on supply of cognate maternal piRNAs, while in the current work differentially supplied maternal piRNAs target a specific region inside of an endogenous cluster. Importantly, maternal piRNAs still targeted the cluster sequences outside of the MiMIC insertion ensuring deposition of H3K9me3 and Rhi. These results suggest that while deposition of H3K9me3 and Rhi to chromatin is directed by maternal piRNAs, they can spread into regions that are not targeted by maternal piRNAs to ensure transcription of piRNA precursors. Similar spreading of Rhi-dependent transcription from piRNA targets to flanking sequences was previously observed for stand-alone TE insertions^2^ and flanks of endogenous piCs^7^. Taken together, our current results and previous observations revealed that maternal piRNAs promote piRNA biogenesis in the progeny by two-prong mechanisms that include activation of non-canonical transcription and post-transcriptional processing in the cytoplasm. Importantly, promotion of transcription can spread into flanking regions, while post-transcriptional processing is local, cannot spread and depends on maternal piRNAs targeting specific genomic regions.

The highly local effect of maternal piRNAs on cytoplasmic processing transforms our understanding of piC function. piCs were considered to be a cohesive unit of piRNA expression with the levels of piRNA output of each cluster proportional to the levels of Rhi-dependent transcription. While our data fully supports this model, they further show that post-transcriptional processing plays a pivotal role in determining the final piRNA output. Importantly, the level of piRNA processing is not uniform along clusters and is determined by the combined levels of *cis-*and *trans-*acting piRNAs targeting specific regions inside the cluster (Fig. 4). The local boost to biogenesis is ensured by ping-pong amplification, which itself depends on the expression of TE targets. Thus, non-uniform processing along piCs likely plays an important role in tuning piRNA biogenesis to ensure repression of active TEs.

### Antagonism between canonical and non-canonical transcription and revision of the ‘transposon trap’ model of adaptation

The piRNA pathway adapts to the fast-evolving repertoire of TEs, however, unlike bacterial CRISPR systems, it lacks machinery to deliberately insert novel sequences into genomic regions encoding RNA guides. Instead, it is believed that the piRNA pathway relies on the activity of TEs themselves: random integrations of TEs into piCs are followed by their repression and by the conversion of TE sequences into piRNA producing loci. Novel TE integrations into piCs were indeed observed after exposure to new TEs in flies and koalas^21,36^. Furthermore, it was shown that insertion of heterologous sequences into piCs generates novel piRNAs from the integrated sequences^19^. On the other hand, studies of adaptation of *Drosophila* strains to new TEs revealed that repression does not immediately follow integration of TEs into piCs^23,37,38^. In addition, unlike TEs which carry regulatory elements that drive their transcription in the germline, previously studied heterologous insertions into piCs lacked promoters, making them inadequate to model processes that occur upon TE integrations.

Here we studied how the activation of canonical transcription from sequences integrated into a piC affect piRNA biogenesis. Surprisingly, we found that a gene integrated into a piC can remain active in the germline and produce a protein product. Furthermore, activation of CT locally disrupts NCT of piRNA precursors and, accordingly, piRNA generation. This process is associated with the removal of Rhi from the region of CT. As Rhi anchors protein complexes required for NCT and export of piRNA precursors^7,8,39,40^ as well as suppression of canonical pre-mRNA processing^8,12,14^, Rhi removal from chromatin might be a direct cause for suppression of piRNA biogenesis. Similar antagonism between CT and NCT was previously reported using deletion of the promoter of a protein-coding gene flanking the *42AB* cluster^7^. Suppression of NCT by CT ensures that protein-coding genes flanking piCs remain active and act as barriers for the distinctive chromatin structure of piCs. In the future, it will be important to investigate the molecular mechanisms of Rhi removal by CT. It should be noted that this process does not require removal of H3K9me3 (Fig. 6A), indicating that other processes beyond binding to this mark are involved in regulating Rhi binding to chromatin.

Our results indicate that genes inserted into piCs can remain active and even interfere with local piRNA biogenesis, suggesting that integration of a TE into a a piC does not guarantee its immediate domestication. Instead, the TE might remain active and counteract piRNA production through its transcriptional activity. How then is a piC-integrated TE eventually repressed and piRNA biogenesis restored? One possibility is that proper domestication requires mutations that disrupt or weaken promoters and regulatory elements of TEs integrated into piCs. In support of this idea, the majority of TE sequences in piCs have accumulated multiple mutations that make them different from the active, consensus sequences. Alternatively, there might be a mechanism that allows the piRNA pathway to overcome canonical transcription to repress TEs and restore piRNA biogenesis. In support of this model, clusters such as *42AB* were reported to harbor several promoters that become active when components of the piRNA machinery such as Mael protein are removed^24^. Taken together, our results suggest that the ‘transposon trap’ model should be revised to incorporate a phase of transcriptional competition initiated by the integration of the new, active TE, before stable silencing is achieved.

## Acknowledgements

We thank Katalin Fejes Tóth and members of the Aravin and Fejes Toth labs for discussion and comments. We thank Julius Brennecke for providing the Rhino antibody. We thank Igor Antoshechkin (Caltech) for helping with sequencing. This work was supported by grants from the National Institutes of Health (R01 GM097363) and by the HHMI Faculty Scholar Award to AAA.

## Materials and Methods

### *Drosophila* stocks

All *Drosophila melanogaster* strains used in this study are listed in Supplementary Table 1. Flies were maintained on standard cornmeal medium at 25°C under standard laboratory conditions.

### Generation of transgenic flies

To generate the UASp-GFP-NLS-SV40 reporter construct, the UASp promoter, GFP-NLS coding sequence, and SV40 polyadenylation signal were amplified by PCR and assembled into the EcoRI/BamHI-digested pBS-KS-attB1-2 vector using Gibson Assembly. The resulting construct was integrated into two genomic landing sites, chr2R:6439020 (BDSC #44828) and chr3L:7575013 (66A6, BDSC #38579)

### RNA HCR-FISH

RNA fluorescence in situ hybridization using hybridization chain reaction (HCR-FISH) was performed with modifications to previously described protocols^41,42^. Probes were designed and synthesized by Molecular Technologies. Fluorophores Alexa Fluor 546, Alexa Fluor 594, and Alexa Fluor 647 were used for signal detection. Images were acquired using a ZEISS LSM880 confocal microscope and processed using ZEN software and Fuji imaging.

### Immunofluorescence microscopy

Immunofluorescence staining was performed as previously described^43,44^. Briefly, approximately five pairs of ovaries were dissected in PBS and fixed in 4% formaldehyde. Samples were washed three times for 10 min in 0.1% Triton X-100 in PBS, mounted in VECTASHIELD antifade mounting medium, and imaged on a ZEISS LSM880 confocal microscope.

### ChIP-seq and ChIP-qPCR

All ChIP experiments were conducted according to a previously established protocol^17^, utilizing anti-H3K9me3 antibody (ab8898), anti-H3K4me2/3 antibody (ab6000) and anti-Rhino antibody (kindly provided by the Brennecke laboratory).

For ChIP-qPCR, immunoprecipitated DNA was quantified using SYBR Green qPCR with MyTaq HS Mix (BioLine). Reactions were performed on a Mastercycler® ep Realplex PCR system (Eppendorf). Ct values were calculated from technical duplicates, and enrichment values were normalized to input DNA and the rp49 control locus. Primer sequences are listed in Supplementary Table 2.

### Small RNA-seq

Total RNA was extracted from dissected ovaries using TRIzol reagent (ThermoFisher #15596018). Approximately 4 µg total RNA was resolved on a 15% polyacrylamide gel, and small RNAs ranging from 19–29 nt were excised and purified. Small RNA libraries were generated using the NEBNext Small RNA Library Prep Kit (NEB #E7330S) and sequenced on an Illumina HiSeq 2500 platform.

### RT-qPCR

Approximately 20 ovaries were dissected and homogenized in 1 mL TRIzol, and total RNA was extracted following the manufacturer’s instructions. RNA was treated with DNase I, and 1 µg total RNA was used for reverse transcription with SuperScript III (Invitrogen).

Quantitative PCR was performed using MyTaq HS Mix (BioLine) with SYBR Green detection on a Mastercycler® ep Realplex PCR system (Eppendorf). Ct values were calculated from technical duplicates, and transcript levels were normalized to rp49 mRNA. Primer sequences are provided in Supplementary Table 2.

### Bioinformatic analysis

ChIP-seq processing and mapping were followed the previous protocol^17^. Sequencing adapters were removed using Trimmomatic (version 0.33)^45^ and cutadapt (version 1.15)^46^, and reads shorter than 50nt were discarded. The first 50 nt of each read were mapped to dm6 genome and vector sequence respectively, using Bowtie^47^ (version 1.0.1, parameters: -v 2 -k 1 -m 1 -t --best -y --strata). Mitochondrial reads were removed prior to downstream analysis. Read coverage and enrichment profiles were generated as RPM-normalized tracks using customized scripts and deepTools^48^. Regions blacklisted by ENCODE^49^ were excluded from enrichment analysis. Read counts across equal-sized bins were calculated using deepTools2 and BEDOPS^50^, and figures were generated using MATLAB. All analysis scripts are available on GitHub: (https://github.com/brianpenghe/Luo_2021_piRNA/blob/main/ChIP-seq.md).

Small RNA-seq analysis followed previously described pipelines^17^. Adapters were removed using Trimmomatic and cutadapt, and reads shorter than 20nt were discarded. Subsequently, reads of specific lengths were extracted: 21-22nt (siRNA), 23-29nt (piRNA), and 21-30nt (small RNA). The selected reads were mapped to the dm6 genome using Bowtie (parameters: -v 0 -a - m 1 -t --best --strata). After removal of mitochondrial reads, read counts were calculated in equal-sized bins using deepTools2 and BEDOPS. Ping-pong signatures were calculated using established method^51^. Phasing analysis was performed as described previously^17,27,28^. Coverage profiles were generated as described in figure legend, and visualizations were produced using MATLAB. Analysis scripts are available at: (https://github.com/brianpenghe/Luo_2021_piRNA/blob/main/piRNA-seq.md).

### *Trans*-piRNA quantification

To identify *cis*- and *trans*-piRNAs with small RNA libraries, we developed an automatic pipeline in Python, called *trans*-piRNA detector (https://github.com/OliveiraDS-hub/trans-piRNA). Following quality control of the small RNA-seq library, we used bowtie v1.2.2^52^ to align the reads to ncRNA sequences from *D. melanogaster* (tRNA, rRNA, miRNA, pre_miRNA, snRNA, and snoRNA) obtained from Flybase r6.60^53^. This first alignment is performed with up to 3 mismatches (parameter -v 3), and unaligned reads with lengths between 23 and 29 bp are extracted as potentially piRNAs (parameter --un). Keeping track of all possible multi-mapper piRNAs in the genome generates heavy and arguably unmanageable file sizes. Therefore, we reduce library complexity by collapsing identical reads with fastx_collapser function from Fastxtools v0.0.14 (http://hannonlab.cshl.edu/fastx_toolkit). The collapsed reads maintain the number of merged reads, allowing quantification of expression afterwards. Then, we use bowtie v.1.2.2 to align the collapsed piRNA-size reads against the *D. melanogaster* genome, obtained from Flybase r6.60. This second alignment allows for 3 mismatches (parameter -v 3), and keeps all multi-mappers (parameter -a). Only reads with at least one alignment in the genome are considered as piRNAs.

The genome position of piRNA clusters for the reference *D. melanogaster* genome was obtained from piRNAdb v2.0^54^. We removed the unistranded clusters *20A* (chrX:21517004-21560994) and *flamenco* (chrX:21631282-21883809) due to the complexity of detecting trans-piRNAs given their particular strandness transcription. piRNAs mapping to clusters are identified with bedtools v2.31.1^55^, function intersect. For each piRNA cluster, we compute *cis-* and *trans-*piRNAs based on different alignment metrics. The metrics are generated based on the total sequences (collapsed identical reads), or total reads (read number in the library). The *cis* quantification, meaning the amount of piRNA produced from each cluster, is measured as “minimum” or “maximum”. In the *cis-*piRNA minimum, we keep only piRNA with 0 mismatches (mm) in the cluster and nowhere else in the genome. This metric differentiates from the classical 0mm unique quantification because it accounts for multi-mappers with 0mm, as long as it is only observed in the same cluster. This technique improves the quantification of *cis-*piRNA because it includes reads from local repeats that may have multiple alignments. The *cis-*piRNA maximum accounts for any 0mm piRNA read mapping to the cluster, regardless of whether they have multiple assignments along the genome, within clusters or not. These two *cis*-piRNA quantifications provide contrasting under- and overestimations of the actual piRNA generated from clusters. Finally, *trans*-piRNA is measured based on piRNA reads mapped with 0mm in a given cluster (unique or multi-mappers), and mapped to other clusters with 1-3mm. The cluster to which a read maps with 0mm is assigned as its cluster of origin, whereas the clusters to which the same read maps with 1-3mm are defined as targets. In our analysis, the cluster-wise quantification of trans-piRNA is based on targeting.

## Data availability

Libraries generated from this study are deposited in GEO under accession codes GSE232906.

## Author Contributions

Y.L. and A.A.A. designed the experiments; Y.L. performed all experiments; D.S.O. performed the analysis to define *cis*- and *trans*-piRNA; Y.L. and P.H. analyzed the data; Y.L., D.S.O. and A.A.A. wrote the paper.

---

**Figure S1, related to Figure 1.**
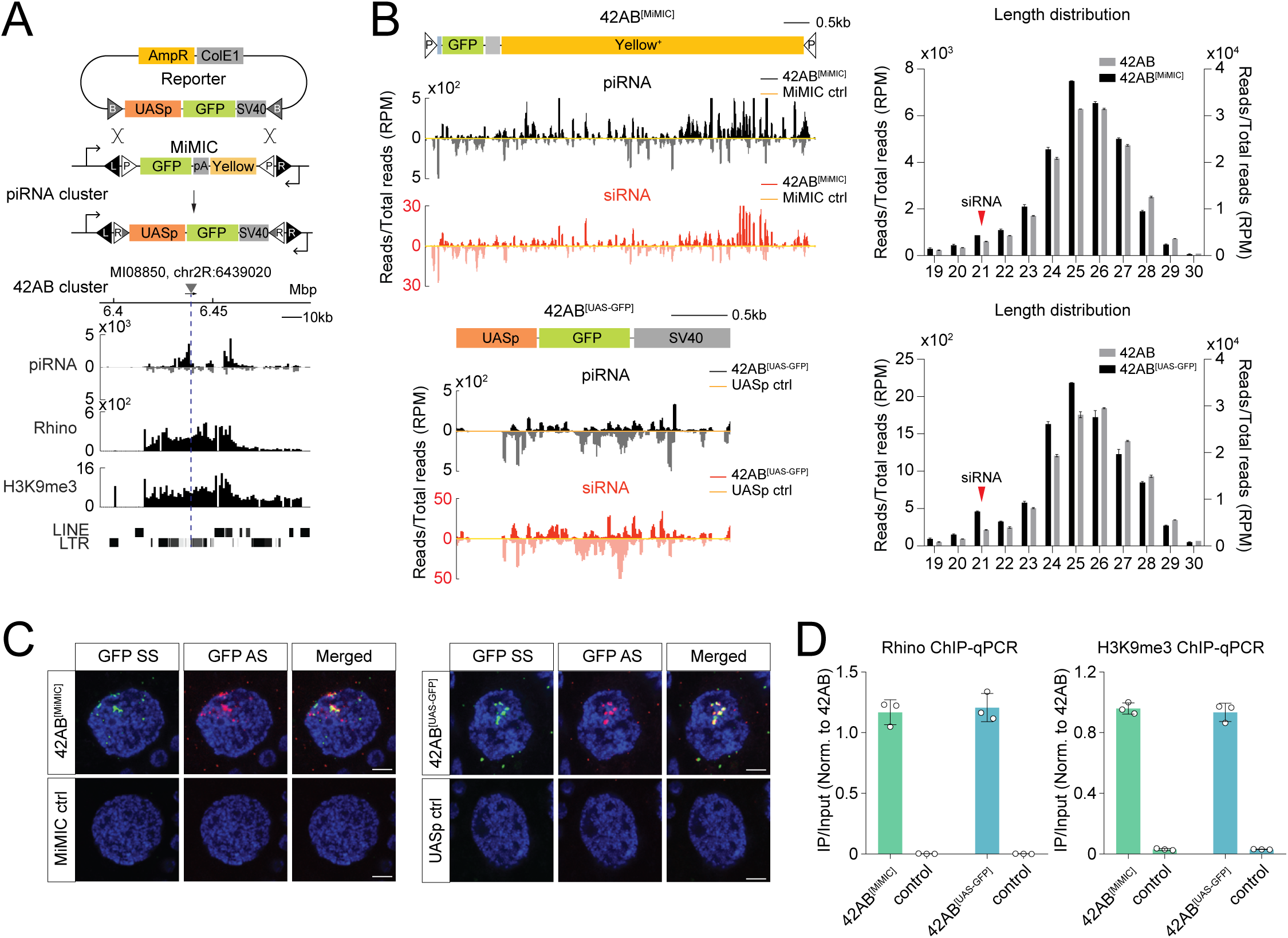
**A. Replacement of MiMIC to UASp-GFP in 42AB cluster.** UASp-GFP reporter was introduced into the *42AB* cluster through recombinase-mediated cassette exchange (RMCE) of the MiMIC insertion. Genome browser tracks show uniquely mapping piRNAs, Rhi ChIP-seq, and H3K9me3 ChIP-seq signals on 42AB cluster. Number of piRNA reads and ChIP-seq reads were normalized to total read counts. **B. Heterologous insertions in 42AB cluster produce piRNA and siRNA.** Shown are profiles of piRNA and siRNA mapping to the MiMIC and UASp-GFP insertions in 42AB cluster. As a control, small RNAs mapping to identical MiMIC and UASp-GFP insertions located in a non-piRNA cluster (non-piC) region are shown. Bar graphs show length profiles of small RNAs mapping to heterologous insertions (black) and *42AB* cluster (grey). Results were normalized to total piRNA reads. Error bars indicate standard deviation of two biological replicates. **C. Detection of nascent transcripts from heterologous insertions.** The nascent transcripts from MiMIC and UASp-GFP insertions in 42AB cluster were detected by strand-specific RNA fluorescent in situ hybridization (FISH) of nurse cell nuclei. Sense and antisense GFP strands are shown in green and red, respectively; DAPI staining of DNA is in blue. Scale bar represents 5μm. **D. Rhi and H3K9me3 mark are enriched on heterologous insertions in 42AB cluster.** Rhi and H3K9me3 ChIP-qPCR experiments were performed on ovaries of flies containing MiMIC and UASp-GFP insertions. Results were measured in three biological replicates and normalized to the region in *42AB* cluster (chr2R: 6,323,128 – 6,323,219). Control is *rp49* gene region. Error bars indicate standard deviation of three biological replicates.

**Figure S2, related to Figure 4.**
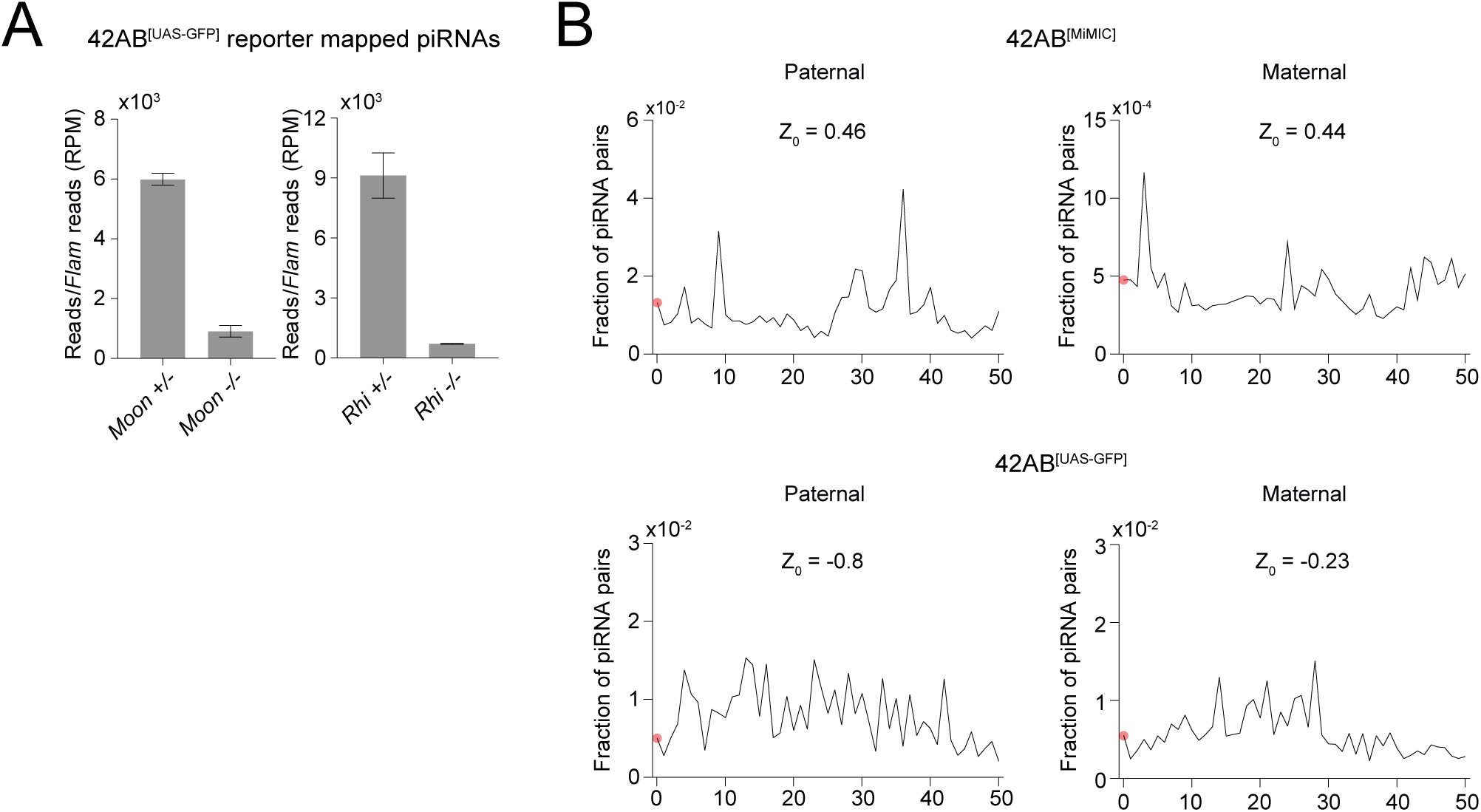
**A. Loss of Rhi and Moon proteins required for NCT leads to reduction of piRNA production from insertion in *42AB* cluster.** The bar graphs show number of piRNA reads mapping to the UASp-GFP insertion in *Moon^FS1Δ1/ FS2Δ28^* and *Rhi^2/KG^* mutants. The reads normalized to piRNA *flam* cluster which is transcribed by canonical machinery and independent of Rhi and Moon. Error bars indicate standard deviation of two biological replicates. **B. The analysis of phasing piRNA biogenesis on MiMIC and UASp-GFP insertions in progenies of maternal and paternal crosses.** No strong phasing signal (Z_0_) was observed in either progeny.

**Figure S3, related to Figure 5.**
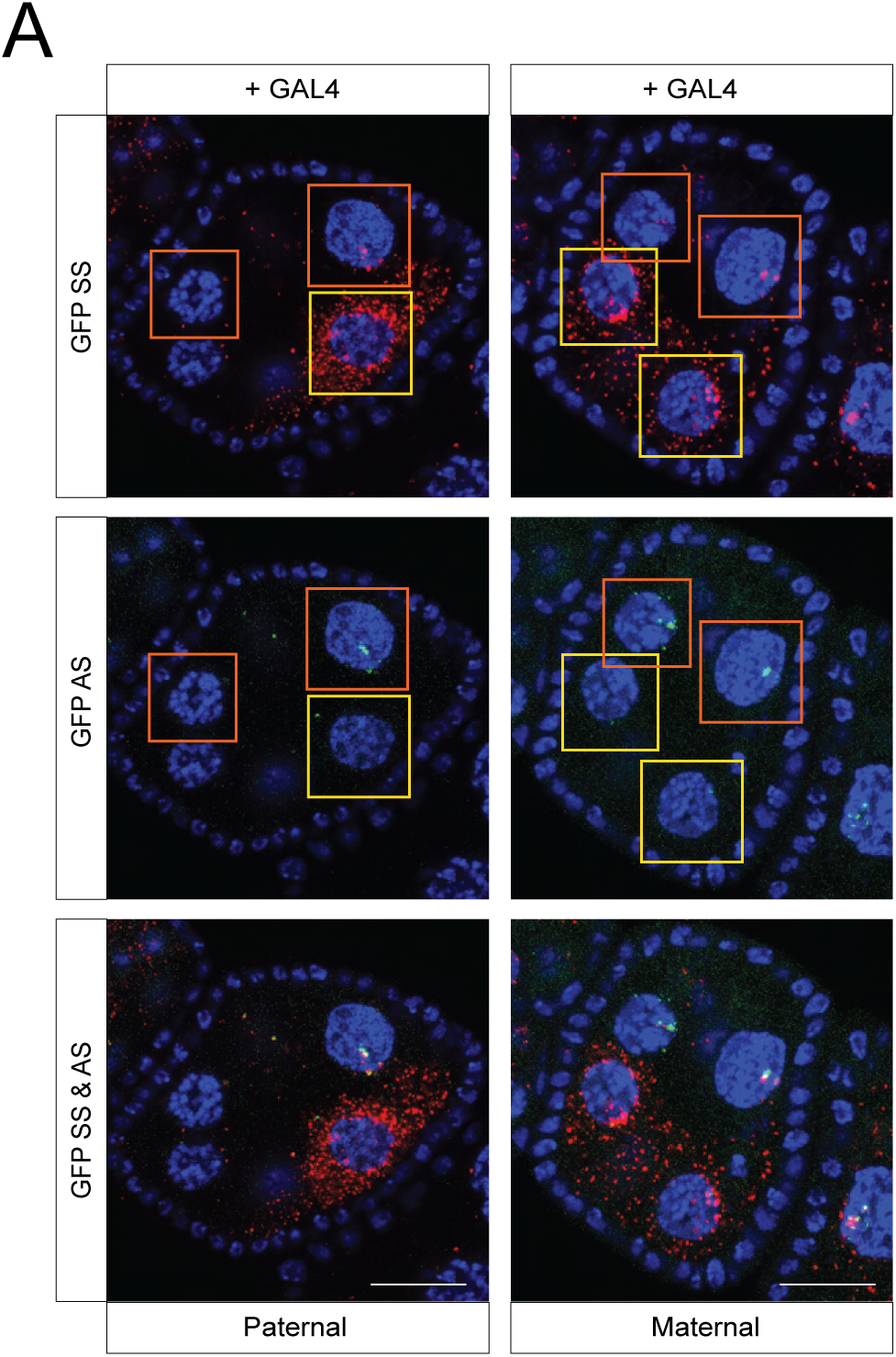
**A. Heterogeneity of UASp-GFP expression upon GAL4 induction between individual cells.** Sense (red) and antisense (green) GFP RNA co-FISH shows nuclei with simultaneous expression of sense and antisense RNA (orange box) as well as cells that have strong expression of sense mRNA and suppression of antisense transcripts (yellow box). Scale bar represents 20μm.

## Notes

### Competing Interest Statement

The authors have declared no competing interest.

